# Songbird parents coordinate offspring provisioning at fine spatio-temporal scales

**DOI:** 10.1101/2022.01.24.477548

**Authors:** Davide Baldan, E. Emiel Van Loon

## Abstract

For parents, rearing offspring together is far from a purely cooperative exercise, as a conflict of interest (‘sexual conflict’) exists over their optimum level of care. Recent theory emphasises that sexual conflict can be evolutionarily resolved, and complete parental cooperation can occur if parents directly respond (‘negotiate’) to each other and coordinate their level of care. Despite numerous experiments show that parents are responsive to each other, we still lack empirical evidence of the behavioural mechanisms by which this negotiation occurs. In this study, we investigated the spatio-temporal coordination of parental provisioning behaviour as a possible mechanism of negotiation over parental care. We deployed an automated radio-tracking technology to track the provisioning activity of wild great tit (*Parus major*) pairs during chick rearing. Our analyses represent the first detailed spatial and temporal description of foraging coordination in songbird parents in a natural context. We demonstrate that the foraging behaviour of the two parents is highly coordinated in space and time, with parents changing their foraging locations in conjunction with their partners’ movements. Therefore, foraging coordination could be a mechanism by which parents directly monitor and respond to each other’s level of investment.

## Introduction

Parental care is a source of both cooperation and conflict for parents (Royle, Smiseth & Kölliker 2012). While caring for young, parents are expected to cooperate as they both invest in the common goal of successfully raising offspring. On the other hand, since parental care is costly for each carer in terms of reduced reproductive opportunities or survival (Williams 1966), each parent is also inclined to exploit the partner and provide a smaller share of the care (hence a ‘sexual conflict’ exists over the evolutionary interests of the two parents) (Trivers 1972; Lessells 2006). A central goal in evolutionary biology is to understand how this sexual conflict is resolved and whether parents can reach a cooperative agreement over how much to care for offspring (Houston & Davies 1985; Lessells 2006; Servedio *et al*. 2019).

Game theoretical models have shown that the evolutionary outcome of sexual conflict depends on the behavioural (‘negotiation’) rules that parents adopt to assess and respond to each other’s level of care over the offspring rearing period (McNamara, Gasson & Houston 1999; McNamara *et al*. 2003; Lessells & McNamara 2012; Johnstone *et al*. 2014; Johnstone & Savage 2019). For instance, early models (McNamara, Gasson & Houston 1999; McNamara *et al*. 2003; Lessells & McNamara 2012) predict that sexual conflict lowers the amount of parental care and reduces parent and offspring fitness compared to a cooperative situation, *i.e*., each carer withholds part of its potential investment to avoid being exploited by the partner (Lessells & McNamara 2012). McNamara, Gasson and Houston (1999)’s and McNamara *et al*. (2003)’s models do not formally specify the negotiation mechanism through which parents monitor the partner’s contribution, but in Lessells & McNamara model, parents decide how much to invest based on the current state of the offspring (which, in turn, reflects the cumulative amount of past investment by the two parents). Recent models, however, show that if parents directly assess each other’s behaviour by coordinating their provisioning activity, such as taking turns of duties, the expected outcome is that parents increase parental care, maximising both parent and offspring fitness (Johnstone *et al*. 2014; Johnstone & Savage 2019) (Fig. 1). Because the evolutionary outcome of sexual conflict strictly depends on how parents acquire information and respond to partner’s care levels, there is a renewed interest to understand the negotiation mechanisms that parents adopt when caring for young (Griffith 2019).

**Figure 1.**
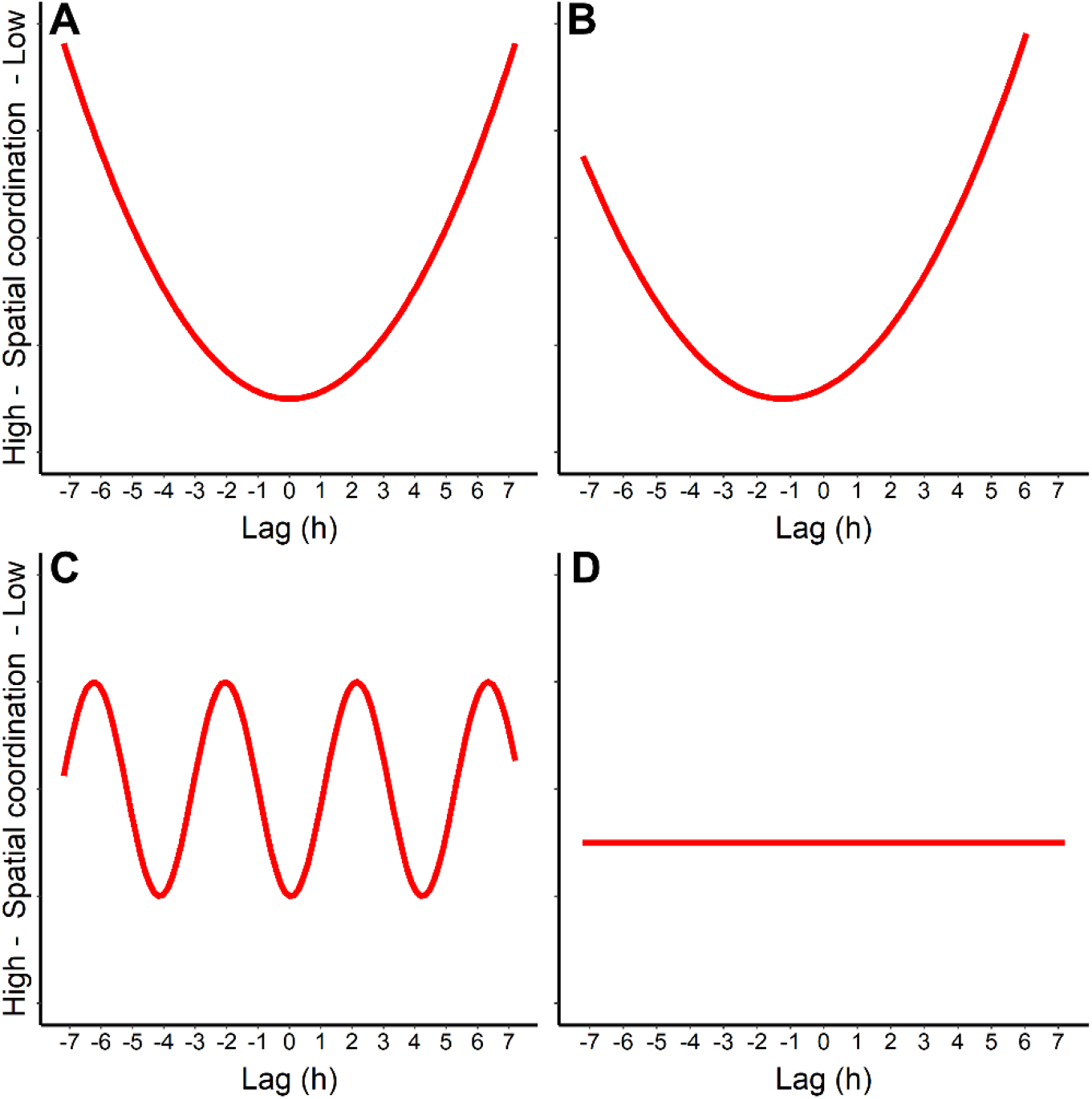
Expected patterns of different forms of spatio-temporal coordination between provisioning parents in the form of correlograms (spatial coordination on the y-axis, and shift in time along the x-axis). **A** occurs when parents coordinate their provisioning in space and time, **B** is the expected pattern if a parent constantly leads the other, **C** represents situations in which parents are coordinated in space and time with a cyclical use of their foraging sites, **D** is the expected pattern when no spatio-temporal coordination occurs, and parents are foraging independently from each other. Note that the direction on the y-axis is flipped for visualization purposes to be consistent with the results.

Negotiation mediated via offspring behaviour has been widely investigated in the field. In birds for instance, parents are highly responsive to offspring begging calls (Kilner & Johnstone 1997) and playback experiments of offspring begging elicit an increase of parental provisioning (Hinde & Kilner 2007). However, it is not yet fully understood how negotiation mediated via direct response to the partner’s behaviour occurs in nature. Empirical studies in songbirds have indicated that individuals modify their provisioning rate in response to the partner’s experimentally manipulated behaviour, e.g., selective playback, handicapping manipulations (reviewed in Harrison *et al*. (2009)). More recently, it has also been argued that parents alternate their visits at the nest more than expected by chance because they actively monitor and respond to each visit of the partner (Johnstone *et al*. 2014; Savage *et al*. 2017); (but see Schlicht *et al*. (2016) and Baldan, Hinde and Lessells (2019)). Therefore, parents are behaviourally responsive to each other, but it is not currently known which behavioural mechanisms are underlying these responses. One possible mechanism is that parents forage in proximity to each other, directly monitor the foraging behaviour of their partner and adjust their own contribution. This knowledge gap hinders our ability to understand which negotiation rules are used by parents while caring the young, and in turn, how sexual conflict could be ultimately resolved (Griffith 2019). And it persists due to the difficulties in collecting provisioning data beyond video recordings or provisioning patterns at the nest (Savage & Hinde 2019).

Fortunately, recent advances in remote monitoring have allowed the study of complex interaction networks between individuals at small spatial and temporal resolution (Krause *et al*. 2013; Smith & Pinter-Wollman 2020). Among these, automated tracking technologies enable researchers to map animal movements and social interplays in detail, especially for species, such as songbirds, that are notoriously difficult to observe in their natural settings due to small size or elevated habitat complexity (Krause, Wilson & Croft 2011; Mennill *et al*. 2012; Snijders, Oers & Naguib 2017). In this study, we deployed an automated radio-tracking technology, Encounternet, to track spatial movements of wild great tit (*Parus major*) pairs during chick rearing with an unprecedented spatio-temporal resolution. We used a metric of similarity in space use between the two parents and analytically shifted (lagged) the movements of one parent relative to the other over time to test the hypothesis that parental provisioning behaviour is coordinated in space and time. If parents forage at different locations over time and are coordinated in space and time, we expect to find the highest parental coordination when time is not lagged and lower levels of coordination when making lagged comparisons (lag zero is the comparison of the location of the two parents at the same whereas lag 1 indicates that spatial locations of one parent is compared with the locations of the other shifted by one time unit, Fig. 1A). If one parent consistently leads the foraging movements over the other, we expect a shift to the side of the leading parent (Fig. 1B). Other non-random scenarios are possible. For instance, if parents are coordinating in space and time but also periodically use their foraging territory, we would expect a cyclical pattern of coordination over time lag (Fig. 1C). Lastly, if parents are foraging independent from each other, we expect coordination not to vary with time lag (Fig. 1D). Using this proposed method of lagging temporal activity and a fine scale monitoring of space use in a free-living songbird, we can have detailed information on the specific behavioural mechanisms underlying parental negotiation.

## Methods

### Study species

The great tit (*Parus major*) is a common passerine species belonging to the *Paridae* family, that easily breeds in nest-boxes throughout Europe, North Africa and Central Asia. It is a model species for studying parental care, negotiation strategies and foraging ecology (Royama 1966; Naef-Daenzer & Keller 1999; Naef-Daenzer, Naef-Daenzer & Nager 2000; Hinde & Kilner 2007). During the chick provisioning period, both parents feed their offspring mostly on Lepidoptera caterpillars, spiders and other insects (*e.g*. adult dipterous insects) with average chick-provisioning visit frequencies up to once every 2-3min per parent (Royama 1966; Naef-Daenzer, Naef-Daenzer & Nager 2000; Baldan *et al*. 2019). Manual radio-tracking studies on chick-provisioning individuals have shown that great tit parents forage by sampling trees nearby the nest site (90% of the foraging locations samples occurred within 45m from the nest in Naef-Daenzer (2000)), searching for available prey (Naef-Daenzer & Keller 1999; Naef-Daenzer 2000). Nevertheless, simultaneous tracking and analysing the movements of both parents has never been conducted.

### Study population and data collection

We conducted this study in 2016 in a great tit population in Roekel, a mixed woodland area in Ede, the Netherlands (52°04’30.7”N, 5°42’48.9”E). This area contains around 250 nest-boxes that we checked weekly from the beginning of April to determine the onset of egg laying and incubation. During early egg laying, we pre-selected active great tit nests to radio-track based on their geographical position. Specifically, we selected nests in which the radio-tracking array would contain relatively homogenous tree coverage in all directions up to *circa* 100 m from the nest site, *i.e*., avoiding nests close to human paths and open fields. In our field site, nest-boxes were positioned every 50 meters and for this study we used nests that were spaced out between 62 and 1125 m from each other.

We caught 16 birds (8 pairs) and fitted these with radio tags during incubation or chick rearing. During the incubation period we caught five males with mist nets nearby the nest, and three females with ‘box nets’ (te Marvelde *et al*. 2011) placed around the nest box. All the remaining individuals were caught and tagged at the nest during chick provisioning. There was no effect of trapping method or timing on brood characteristics such as hatching date, brood size or parental behaviour such as provisioning rate during data collection (Mann-Whitney tests, all *P* > 0.3).

We collected radio tracking data with the automatic tracking system Encounternet (Encounternet LLC, Portland, OR, U.S.A.). Encounternet consists of small radio transmitters of 0.9 g (5% of the body mass of our studied individuals), fitted to the bird with a leg-looped backpack harness (Rappole & Tipton 1991). These tags broadcast a radio signal every 5s, which is recorded by small wireless receivers logging the ID number, time and received signal strength indication (RSSI) of every tag pulse they receive (Mennill *et al*. 2012). To track spatial movements of the eight tagged pairs during chick provisioning, for each pair we deployed 37 receivers around the nest site in a 75m array. We placed these receivers in a triangular array consisting of three ‘rings’ at 25, 50 and 75 meters from the nest (Fig. 2). We positioned the receivers in trees or plastic poles at *ca*. 3.5m height at a regular distance of approximately 25m distance from one another. We located and surveyed the coordinates of the receivers in the field with a survey-grade GPS (Ashtech ProMark 800, Santa Clara, CA, U.S.A.). At the nest site, we placed one more receiver on the front side of the tree *ca*. 50 cm above the nest-box. On average, parents spent 73% of their time within the array detection area (mean ± SE: 0.73 ± 0.05) and the proportion of time inside of the array negatively correlated with the number of chicks (GLMM, estimate ± SE: −1.12 ± 0.25, χ^2^ = 9.86, d.f. = 1, *P* = 0.002, N = 128). In addition to the Encounternet array, a small video camera was mounted in the roof of the nest-box and connected to an external video recorder at the foot of the tree. Video recordings (720 x 576 pixels of resolution) started before 0730h and ended on the recording days and were synchronized (to the nearest second) to the Encounternet array to simultaneously monitor spatial movements and visits at the nest of the provisioning parents. We positioned the Encounternet array and the video set-up the day prior to data collection to habituate parents to their presence. We tagged all the parents at least two days before data collection to reduce possible effects of tagging on provisioning activity of the parents.

**Figure 2.**
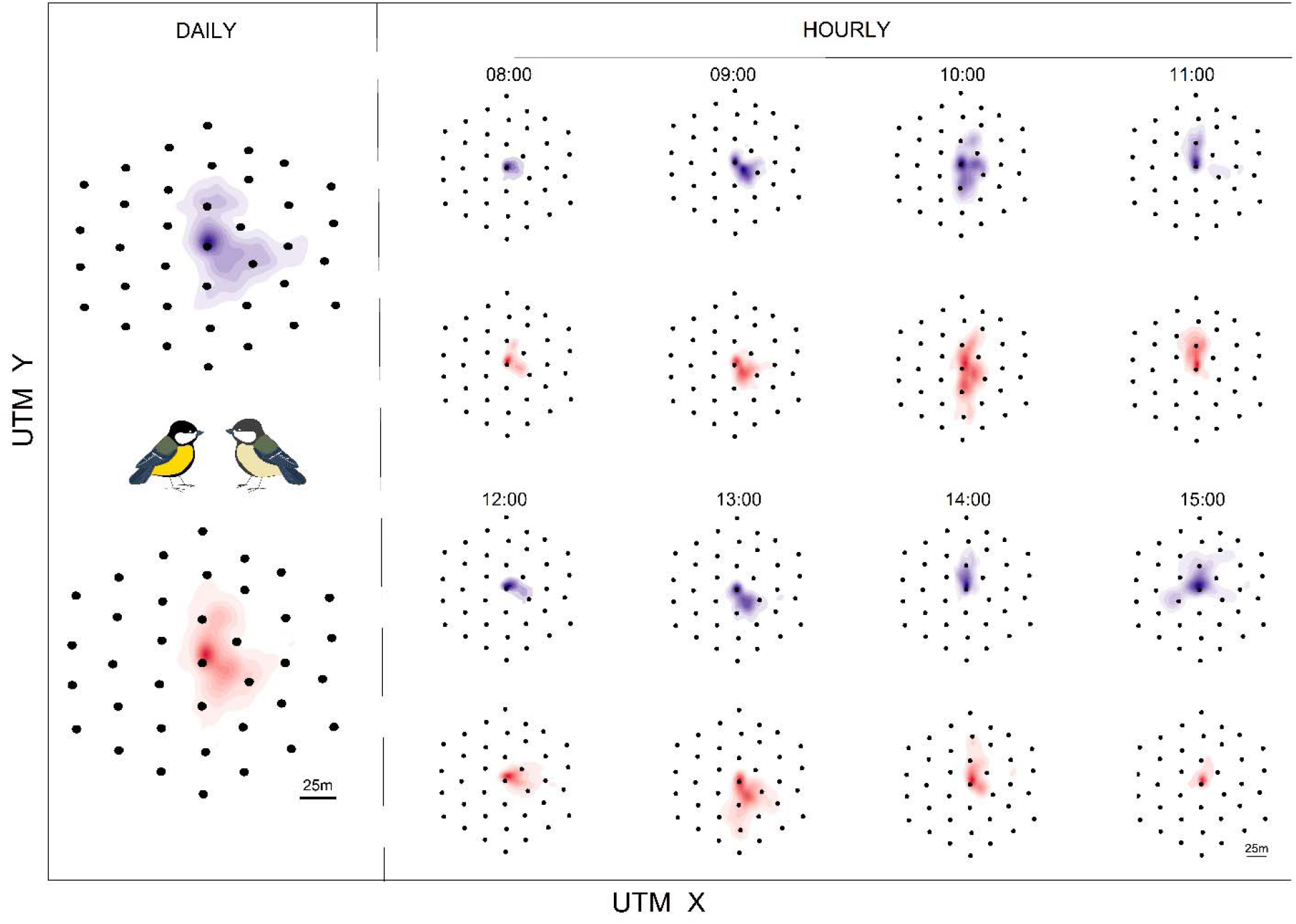
Example of utilization distributions (UDs; space use) of a single pair (male and female, respectively in blue and red). The two leftmost UDs represent male and female daily space use over the full period of eight hours. The other 16 UDs, represent the hourly UDs for each sex. UTM represents the geographical coordinates of the Encounternet array expressed as Universal Transverse Mercator such that one unit corresponds to one meter. The black dots are the Encounternet receivers used to automatically track pairs in their environment with the centre being the nest site.

For each Encounternet nest, we collected radio tracking data of both parents for four consecutive days as part of a brood-size experiment. Here, we used 64 hours of data (eight hours from 0800h to 1600h for all nests) collected on the first day of the four-day period under natural and unmanipulated conditions. From the video recordings, we detected a total of 1783 provisioning trips for which we scored the parental sex (determined from the blackness of the crown feathers) and the times that the bird entered and left the nest-box (to the nearest second). We used a triangulation algorithm implemented in MATLAB (The MathWorks, Natick, MA, U.S.A.) to locate the position inside the array of each tagged parents every five seconds from the radio signals logged by the receivers. This triangulation algorithm provides estimated locations with an accuracy of 13.62 ± 0.54 m (mean ± SE) (for further details on the triangulation algorithm and validation test see supplementary material S1). We estimated a total of 6753 unique locations within the 64 hours used in this study. At the end of the four-day study period, we removed the radio-tags from the parents by catching them at the nest. Our birds were equipped with tags for an average of 10.5 days (range 6-26 days).

### Calculation of similarity in spatial utilization distribution between parents

To investigate spatial coordination between provisioning parents, we grouped the location-data per hour and applied the dynamic Brownian bridge movement model (Kranstauber *et al*. 2012) to estimate the utilization distributions (UDs) of each parent per hour (Fig. 2). This method uses the time series of sequential locations for each individual and summarizes its movement into a 2-dimensional spatial representation referred to as utilization distribution (Worton 1989). A comparison between UDs of different individuals, via indices of spatio-temporal overlap (Fieberg & Kochanny 2005), have been used to explicitly quantify interactions between individuals (Robert, Garant & Pelletier 2012; Schauber *et al*. 2015; Lewis *et al*. 2017). In this study, we compared male and female UDs by using the earth mover’s distance (EMD), a measure that quantifies the similarity between two UDs (Kranstauber, Smolla & Safi 2017). Note that EMD increases with increasing dissimilarity between two UDs, whereas it assumes a value of 0 when two identical UDs are compared (Kranstauber, Smolla & Safi 2017). We used EMD as measure of spatial coordination (similarity) of male and female parents, with a high EMD value representing low spatial coordination. Calculations of the parental UDs via dynamic Brownian bridge movement models and EMD were performed in the R package *‘move’* (Kranstauber, Smolla & Scharf 2017).

### Statistical analysis

To investigate whether parents coordinate foraging in space and time we carried out analyses at different temporal scales: a broad scale (among hours) and a fine scale (among 10 minute intervals) analysis. In the broad scale analysis, we used male and female hourly utilization distributions (UDs). For each nest and each day, we calculated the earth mover’s distance (EMD) between the male and female UDs of the same one-hour period (e.g. male UD from 0800h to 0900h with female UD from 0800h to 0900h). These EMD values were defined as “lag 0” EMD as they represent the observed similarity in the parental space use over the same time period. To explore whether parents are coordinated in space and time, within each nest and day, we calculated the EMD between all combinations of male and female hourly UDs, by progressively lagging the UDs of one parent relative to the other. For instance, by lagging the female UDs by one hour, we compared the EMD between male UDs from 0900h to 1000h with female UD from 0800h to 0900h and so on. These EMD values are defined as “lagged” EMDs, as they represent the similarity in the parental space use when one individual provisioning activity is lagged relative to the other. In this way, we expected that if parents are coordinated in space but also in time, the EMDs at lag 0 would be smaller (higher similarity between the UDs) compared to the lagged values in absence of any periodicity in foraging territory use (Fig. 1A). On the other hand, if parents are foraging independently from each other at different locations over the course of the day, we would expect the EMD at lag 0 not to differ from the lagged values. We would also expect this pattern (the EMD at lag 0 not to differ from the lagged values) if both parents would constantly forage at the same location over the day, regardless of the underlying mechanism (coordination or independent movement). To test for spatio-temporal coordination between parents, we investigated whether the EMD value of a parental UD (focal UD) was smaller when matched with its own partner’s UD at the same time (lag 0) or with its own partner’s UD at different time (lagged EMD). This approach created a pseudo-replication issue since the same UD was present multiple times in the dataset. To resolve this problem, this analysis was performed using linear mixed models with EMD as response variables, “lag” as fixed effect, and “male UD ID” and “female UD ID” nested into “Nest ID” as random factors. The variable “lag” was used as factor in the analysis. We also ran post-hoc tests to investigate the differences in EMD between the different lag classes. We carried out this broad scale analyses based on 1-hr intervals to broadly investigate whether great tit parents forage together over multiple locations over the course of a day and to rule out the possibility that the observed level of coordination is solely a by-product of parents using the same foraging patch simultaneously but independently from the partner.

Subsequently, we investigated spatio-temporal coordination between parents at a finer scale by decomposing parental hourly UDs into six UDs, each covering a ten-minute interval. In this fine scale analysis, we applied the same methodology and statistical models of the broad scale analysis (in the fine scale analysis the lag between UDs occurred in steps of 10 min periods instead of one-hour periods) to explore whether pair coordination in space also occurred at a smaller temporal scale. We chose the 10-minute interval as unit of analysis because great tit foraging trips usually have a duration of few minutes (one visit every 3.76 minutes in this dataset). Our 10-minute blocks enable us to have a more precise indication of the temporal scale to which parents are responsive to changes of partner’s movement while having a sufficient number of locations (120 locations) to reliably generate parental UDs.

In addition to the previous analysis, we explored whether pair coordination varied over time (such as time of day, period during breeding season) or was related to brood size and age. For this we used linear mixed models with EMD as the response variable and ‘hour of the day’, ‘day in April’, ‘number of chicks’ and ‘chick age’ as fixed effects. The correlation coefficients among these variables were small (R^2^ < 0.3), so that they could be included in the same model without problems.

To better explore the extent to which foraging movements are coordinated between the parents we also carried out an analysis of the foraging angles and an analysis of proximity between the parents. For the analysis of the foraging angles, we calculated the angles of each parental location relative to the nest site and correlated males’ and females’ angles at each time point. Here we used a Circular correlation test (function *cor.circular)* implemented in the R package *circular* (Agostinelli & Lund 2017). Because angles of consecutive locations are highly correlated (Pearson correlation of 0.72 for a lag of 5s =; and 0.53 for a lag of 60s), we carried out the same correlation test on a resampled subset of the data by using one location every minute to not inflate the statistical results. For the analyses of proximity, we defined the two parents to occur in proximity with each other when they were located within 10m at the same time point (as in Snijders *et al*. (2014)). As the Encounternet accuracy in this study is 13.6m, we consider our 10m cut-off to be a conservative value to estimate parents’ encounters. In this way, we explored the locations in which parents were found in proximity and grouped them by binning distances every 10m from the nest site until the edge of the array. In this way, we investigated in which areas parents are more frequently in contact with each other. We did this by fitting a linear mixed model with ‘cumulative time spent in proximity’ as the response variable, ‘distance from the nest’ (treated as categorical variable) as fixed effect and ‘Nest ID’ as random factor. Lastly, we explored whether parental coordination of the provisioning was related to period of higher provisioning activity. Here we correlated the hourly values of EMD as measure of foraging coordination with the hourly provisioning rate at the nest.

All the statistical analyses were performed in R environment (version 3.2.3). All mixed models were performed with the *lme4* package (Bates *et al*. 2015). We used a backward selection procedure, starting with the full models containing all the main effects, then dropped the predictor with the highest P-value in each step until only significant effects remained in the final model if any. The significance of the main effects was calculated with the Kenward-Roger approximation implemented in the *pbkrtest* package (Halekoh & Hojsgaard 2014). In all models, the proportion of available locations for each period used to generate the UDs was included to weight the cases.

## Results

In the broad scale analysis, earth mover’s distance (EMD) between male and female UDs differed between lags (F_14,231_ = 2.71, *P* = 0.001; Fig. 3A). Post hoc comparisons showed that EMD was significantly smaller at lag 0, indicating that parental provisioning activity was coordinated in space and time (Fig. 3A). This pattern was not influenced by the presence of un-estimated locations produced by parents foraging outside the detection zone of the Encounternet array (see supplementary material S2). The same pattern was found in the fine scale analysis (F_10,1765_ = 21.87, *P* < 0.001, Fig. 3B) showing that parental coordination occurred at a short temporal scale (ten minutes). Hour of day, day in April, number of chicks and chick age did not influence parental coordination (Table 1).

**Figure 3.**
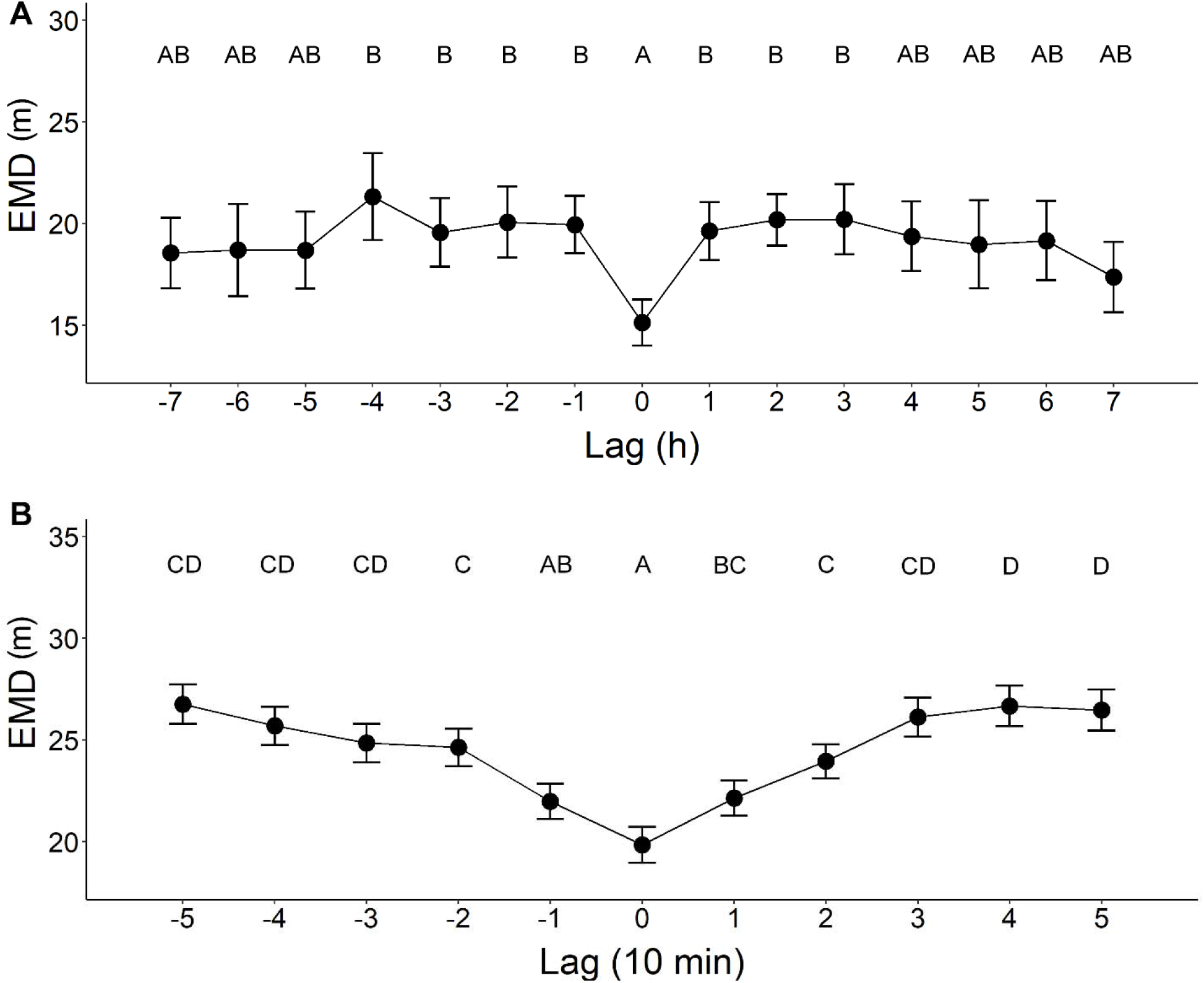
Broad **(A)** and fine **(B)** scale analysis of similarity in parental spatial utilization distribution (EMD) in relation to lag. **A** – Effect of lag on EMD in the broad scale analysis. Positive lag values refer to a situation when female UDs are compared with male UDs at an earlier point in time (e.g. female UD at 1000h with male UD at 0800 has a +2 lag). Negative values of lag refer to the opposite situation when male UDs are compared with female UDs at an earlier point in time (e.g. male UD at 1000h with female UD at 0800 has a lag value of −2). **B** – Effect of lag on EMD in the fine scale analysis. Also in here, positive values of lag occur when female UDs are compared with male UDs occurred earlier in time, whereas negative values of lag occur when male UDs are compared with female UDs occurred earlier in time. Mean ± SE are shown in the graphs. Different letters on top of the datapoints indicate significant differences among lag classes in the post-hoc tests.

**Table 1.** Estimated parameters of the linear mixed models investigating the effect of nest characteristics on similarity in parental space use (EMD). ‘Nest ID’ was included as random effect in the models. For each variable, the statistics expresses whether the model including this variable is explaining more variance than the smaller model, without the respective variable.

| Variables | F-test | ndf | ddf | P-value |
| --- | --- | --- | --- | --- |
| Hour of day | 0.39 | 1 | 31.17 | 0.536 |
| Day in April | 4.11 | 1 | 5.31 | 0.095 |
| Number of chicks | 0.09 | 1 | 3.14 | 0.774 |
| Chick age | 1.68 | 1 | 3.60 | 0.271 |

The angle of the foraging locations significantly correlated between the two parents (*r* = 0.38; *t* = 40.07, *P* < 0.001 for the full dataset; *r* = 0.37, *t* = 13.58, *P* < 0.001 for the restricted dataset) revealing that great tit parents foraged together and visited together multiple locations in their territories (Fig. 4A). In their foraging activity, parents were on average at a distance of 34.7m from each other (range: 0.29-123.29m; Fig. 4B). Great tit parents occurred in proximity with each other both at the nest site (within 10m from the nest location) and at the foraging areas (Fig. 4C). In particular, the frequency of proximity significantly differed between areas (F_6,42_ = 4.87, *P* < 0.001). Lastly, we found a significant correlation between EMD and provisioning rate (F_1,30_ = 4.98, *P* = 0.03; Fig. 4D); hourly periods of higher provisioning at the nest were associated with greater spatial coordination between the two parents (indicated by lower EMD values).

**Figure 4.**
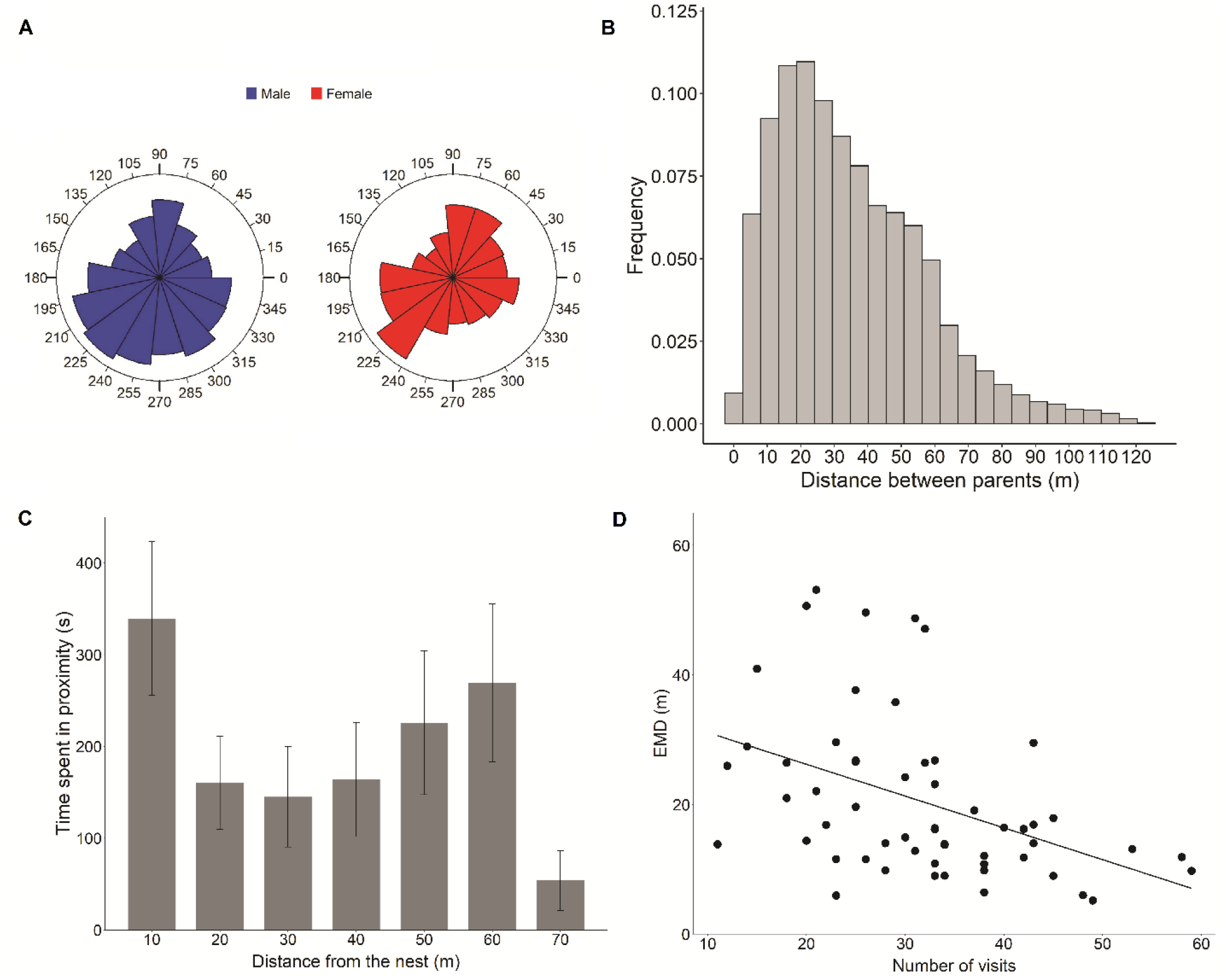
**A** – Histogram of the angles of the foraging locations for male (in blue) and female (in red) great tit parents. **B** – Frequency histogram of the distances (in meters) between male and female parents. **C** – Cumulative time in which male and female parental locations occurred within 10m from each other at different distances from the nest. Mean ± SE are shown in the graph. **D** – Relationship between similarity in parental space use (EMD) and number of visits at the nest during 1-hour periods.

## Discussion

We deployed an automated radio-tracking system to investigate whether great tit parents coordinate their provisioning movements in space and time, as measured by similarity in parental space-use. By using a methodological approach in which we analytically alter the temporal activity of one parent over the other, we showed that spatial coordination is higher when the activity of one parent was not lagged over time, demonstrating that the movements of foraging great tit pairs are highly coordinated in space and time. In addition, parental encounters (periods in which parents occurred in close proximity with each other) occurred both at the nest site and at the foraging locations, and parents’ level of coordination positively correlated with the rate of chick provisioning. These results provide concrete evidence of spatio-temporal coordination of parental provisioning in wild songbirds.

To our knowledge, this is the first time that fine scale spatial data has been collected on pair coordination in songbird parents in a natural setting. In zebra finches (*Taeniopygia guttata*), Mariette and Griffith (2015) found that pairs coordinated their foraging by synchronizing their visits to feeders deployed in their territory. Our study reinforces their findings, showing that pair coordination also occurs in natural situations where individuals were free to search for food in their environment. Moreover, by analytically lagging the provisioning activity of one parent over the other by blocks of ten minutes, we show that this spatio-temporal coordination takes place on the order of a few minutes in natural conditions. Because foraging trips also occur at a similar time scale (in this dataset parents visits on average every two minutes), these findings indicate that parents can continuously monitor and promptly respond to the spatial movements of the partner. Our study provides one possible negotiation mechanism for decades of studies that have shown parental behavioural adjustments due to experimental manipulation (see Harrison *et al*. (2009) for a meta-analysis on the topic). In conjunction with spatial coordination of foraging, another non-mutually exclusive mechanism of negotiation is vocal communication within the pair (Mariette 2019). In birds, parents often vocally communicate with each other to coordinate activities, such as incubation breaks (Boucaud *et al*. 2017), so coordinated spatial movements between parents could also be regulated by contact calls or calls conveying information on newly discovered food sources. Future studies should combine spatial and acoustic data to reach a comprehensive understanding of the negotiation mechanisms in use during parental care.

While movement data on foraging parents are common and available in other bird species, such as sea birds and raptors (Cagnacci *et al*. 2010), studies on parental coordination in those species are recent (Tyson *et al*. 2017; Wojczulanis-Jakubas, Araya-Salas & Jakubas 2018; Grissot *et al*. 2019; Kavelaars *et al*. 2021). For pelagic species in particular, the attention has been primarily directed to the study of dual-foraging strategy, in which alternation of short, chick-provisioning and long, self-provisioning foraging trips by the two parents occurs as a mechanism to steadily deliver food over time to the offspring (Shoji *et al*. 2015; Tyson *et al*. 2017). Nevertheless, these studies have not formally quantified similarity in parental movements and have not proposed a concrete explanation for how parents gain information on the partner’s activity in the context of negotiation rules (but see Kavelaars *et al*. (2021)). In this study, we investigate fine scale coordination of the foraging trips and propose that constant spatial proximity with the partner could function as a monitoring mechanism of the mate’s investments.

Our radio-tracking data and analysis of the foraging angles revealed that great tit parents forage together and sample multiple locations in their territory over the course of the day. These findings confirm previous radio-tracking studies on single great tit individuals showing that provisioning individuals changed foraging locations over time when a food patch become unprofitable (Naef-Daenzer & Keller 1999; Naef-Daenzer 2000). In our study, we did not quantify and measured food availability and distribution as in Naef-Daenzer (2000), and therefore we have no information on parental decisions to change foraging patches. However, by finding that parents switched foraging locations over time and by showing that a change in the foraging locations by one parent is matched by a change in the partner’s movements in the same direction within a ten-minute interval, we showed that foraging decisions also depend on the behaviour of the breeding partner. The only other possible explanation for these coordinated movements, although we consider it unlikely given the temporal scale examined in this study, that does not involve any form of interaction/information exchange between the parents, is that both parents, simultaneously and independently from each other, constantly are aware and move to newly emerged food patches during the day. We also found that parents forage closer to the nest (within the array) when there are fewer chicks to provision and parental coordination was higher at high feeding rates. This relationship between home range size and offspring number could be created by resource depletion and low renewal rate of food items in a patchy environment (Ford 1983). In our provisioning great tits, parents feeding larger broods may have already depleted food sources closer to the nest by the time our data were collected (between day nine and 13 of chick age), and therefore had to forage further away from the nest. Furthermore, in periods of more intense foraging activity parents may benefit from foraging together by promoting information exchange about patch availability (Valone 1989). Taken together, these findings suggest that foraging decisions of chick-provisioning parents may be regulated both by their need to monitor (and negotiate with) their partner but also to maximize energy delivered to the offspring (Orians & Pearson 1979; Olsson, Brown & Helf 2008). Future studies should integrate radio-tracking of foraging pairs and food distribution data at fine spatial scales, e.g. via frass sampling or branch samples (Zandt 1994; Naef-Daenzer & Keller 1999), to integrate in a comprehensive framework both negotiation and central place foraging rules during the chick provisioning period. Other than being a potential negotiation and foraging mechanism, parental coordination could also act as an antipredator strategy. Social foraging has been shown to decrease predation risk via diluted pre-capita risk of being predated or more efficient vigilance (Caraco 1981; Wrona 1991; Sorato *et al*. 2012). Therefore, it is possible that coordinated movements between parents could have multiple functions for the breeding pairs during the chick provisioning period.

The finding that parents coordinate their provisioning activities in space and time has important evolutionary implications in the resolution of sexual conflict. The two most recent theoretical models have emphasized that if parents negotiate by directly monitoring and responding to each other, they can maximize their investment in care, leading to complete cooperation and higher parental and offspring fitness (Johnstone *et al*. 2014; Johnstone & Savage 2019). Our study provides evidence of a mechanism, spatio-temporal coordination of the provisioning, by which parents negotiate and respond to partners’ activity. This is a first step in our understanding of how the specific rules that parents use during care affect their investment decisions and ultimately how sexual conflict is resolved withing pairs (Lessells & McNamara 2012; Johnstone *et al*. 2014; Griffith 2019; Servedio *et al*. 2019). Because the great tit pairs used in this study were also part of a later brood size manipulation experiment and do not belong to a long-term monitored study site (Baldan *et al*. 2019), we could not reliably investigate fitness consequences of coordinated provisioning nor its relationship with pair bonding behaviour (Griffith 2019; Culina, Firth & Hinde 2020). However, experimental manipulations of parental coordination, *i.e*., via selective feeders that allow only specific individuals to gain access to prey items (Aplin *et al*. 2015; Sonnenberg *et al*. 2019), will be needed to further test the causal link that coordination ameliorates sexual conflict and increases offspring fitness.

Understanding how cooperation between parents can evolve and what are the proximate mechanisms are fundamental questions that have recently seen a growing interest (Griffith 2019; Servedio *et al*. 2019). In this study, we demonstrate the existence of fine scale coordination of offspring provisioning activity by parents, which has been proposed to promote cooperation between breeding pairs and ameliorate sexual conflict. Uncovering the physiological and genetic mechanisms underlying variation in coordination (Donaldson & Young 2008; Taborsky & Taborsky 2015; Fischer, Nowicki & O’Connell 2019) and possible environmental constraints on the emergence of coordinated care (Baldan & Ouyang 2020) will also be necessary to reach a comprehensive understanding of the evolution of parental care.

## Acknowledgements

We thank the Municipality of Ede for the use of their terrain and facilitating our research. We are grateful to Peter de Vries, Salome van ⍰t Riet and Mathias Cox for their help in the field. We thank Catherine M. Lessells for conception and design of the project, as well as her comments on the manuscript. We also thank Marleen Cobben, Kamiel Spoelstra, John Burt, Victor de Jager, Lysanne Snijders and Marc Naguib for their help on MATLAB and Encounternet algorithm, and Marcel Visser, Arie van Noordwijk and Jenny Ouyang for discussion and comments at different stages of the manuscript preparation. We thank the members of the UNR Evoldoers lab group for comments on earlier drafts of this manuscript. We are grateful to the editor and three anonymous reviewers for helpful comments.

## Author Contributions

D.B and E.E.v.L conceived the study. D.B. collected the data. D.B. and E.E.v.L analysed the data and wrote the paper. All authors revised, edited and approved the manuscript before submission.

## Data availability statement

Data available from the Zenodo Repository: https://doi.org/10.5281/zenodo.4722817.

## Ethics statement

Permission for this study was granted by the Dutch legal entity: KNAW Dier Experimenten Commissie (DEC) no. NIOO-14.17.

## Declaration of Interests

The authors declare no competing interests.

## Funding

This research was supported by a grant (823.01.005) from the Netherlands Organization of Scientific Research (NWO) to Catherine M. Lessells.

## Notes

https://doi.org/10.5281/zenodo.4722817

### Competing Interest Statement

The authors have declared no competing interest.

